# miR-425 suppresses EMT and inhibits the development of TNBC (triple-negative breast cancer) by targeting TGF-β 1/SMAD 3 signaling pathway

**DOI:** 10.1101/477877

**Authors:** Yingping Liu, Hongfei Qiao, Jinglong Chen

## Abstract

**Background:** EMT has the crucial effect on the progression and metastasis of tumor. This work will elucidate the role of miR-425 in EMT and development of TNBC.

**Methods:** The differential miRNA expression among non-tumor, para-tumor (adjacent tissue of tumor) and tumor tissues was analyzed. The luciferase activities of TGF-β1 3’ UTR treated with miR-425 were determined. Then human breast cancer cell lines were dealt with mimics or inhibitors of miR-425, and then the cell proliferation and migration, invasion ability were assessed. The expression of TGF-β1 and markers of epithelial cell and mesenchymal cell were analyzed. The influences of miR-425 on development of TNBC through inducing EMT by targeting TGF-β 1 and TGF-β1/SMAD3 signaling pathway in TNBC cell lines were investigated. Furthermore, Xenograft mice were used to explore the potential roles of miR-425 on EMT and development of TNBC *in vivo*.

**Results:** Compared with non-tumor tissues, 9 miRNAs were upregulated and 3 miRNAs were down-regulated in tumor tissues. The relative expression of miR-425 in tumor tissues was obviously much lower than that in para-tumor and non-tumor tissues. MiR-425 suppressed TGF-β1 expression, additionally inhibited expression of mesenchymal cell markers, while exerted effects on cell proliferation and migration of TNBC cell lines. Moreover, the agomir of miR-425 could protect against development process in murine TNBC xenogarft model.

**Conclusions:** Our results demonstrated that miR-425 targets to TGF-β1, and was a crucial suppressor on EMT and development of TNBC through inhibiting TGF-β1/SMAD3 signaling pathway. It suggested that aim at TGF-β1/SMAD3 signaling pathway by enhancing relative miR-425 expression, was a feasible therapy strategy for TNBC.

## Introduction

In worldwide, the most usual cancer of women is BC (Breast cancer) [1]. Moreover, the morbidity of BC is the seventh among all cancers, and its mortality rate is third in the women world [2]. According to different prognoses and options of treatment, BC is divided into several subtypes disease associated with heterogeneity [3, 4]. In clinical, corresponding to expression of ERBB2/HER2 (epidermal growth factor receptor 2), PR (progesterone receptor), and ER (estrogen receptor), patient with BC are classified as three subtypes of triple-negative, HER2+, and ER+/PR+. The overall prognosis of patients with Her2+ or ER+/PR+ subtypes has been dramatic improved by trentment with HER2 targeted hormone therapy. But, base on the aggressiveness of TNBC (triple-negative breast cancer) and the lack of effective targeted therapy, the overall survival rate of patients with TNBC is the worst [4]. About 20% patients with BC is TNBC, and the high metastasis rate could be seen in patients with TNBC, such as visceral organs (brain, liver, and lung, etc.) [5]. Therefore, there is an urgent need to develop new therapeutic interventions for preventing and treating metastasis of TNBC.

EMT (Epithelial to mesenchymal transition) is the critical course, and has the important role in metastasis and progression of cancer. The invasion and migration abilities increase, and epithelial polarity vanishes away during this process together with epithelial markers down-expression and mesenchymal markers over-expression [6-13]. Some transcription factors regulate the process of EMT, including Zeb1, Slug, Snail, TGFβ, and Twist [9, 13]. The biological process of EMT is complex, including losing characteristics of epithelia, and acquiring mesenchymal phenotypes. Moreover, the expressions of mesenchymal cell markers, such as Smad 3, α-SMA (alpha smooth muscle actin), etc, were promoted [14, 15]. The growth of tumor relies not only on tumor malignancy but also the relationship between stromal and tumor cells [14, 15]. The system of tumor vessel includes pericytes, smooth muscle cells and, epithelial cells (ECs), which plays an important role in tumors progression and angiogenesis activation [16, 17]. The EMT process of ECs is conduced to the progression of cancer, that has been identified by recent researches [18-20]. In this process, the properties of migration or invasion and markers of mesenchyma (e.g., α-SMA, FSP-1) are obtained, the epithelial markers (e.g., CD31, VE-cadherin) of ECs and junctions of cell-cell were losed [21]. These investigations confirm that all kinds of pathological processes occurs with, for example, cancer [18, 19, 22]. Therefore, inhibition of EMT process may be a good tumor treatment strategy.

Lately, some scholars have revealed that TGF-β1 is a cytokine with multifunction, and plays a crucial role in pathophysiology of BC [23]. The canonical and noncanonical TGF-β1 signaling pathways gene mutations encoding TGF-β1 or the change of TGF-β1 receptors associates with BC oncogenic activity. In BC, the progression of epithelial cell cycle is inhibited by TGF-β1 at early phase while apoptosis is promoted, which indicates its suppressive effects on tumor [24]. But, this cytokine promotes tumor progression in the late phase and associated with metastasis or invasiveness of cancer, and higher motility of cancer cell. In addition, TGF-β1 induces EMT, and is related to modification of microenvironment in cancer [25]. The well known "TGF-β1 paradox" is called to describe this phenomenon, which suggests TGF-β1 can be used for the novel therapy approaches on BC with better understanding its functions [26]. By protecting against TGF-β1, several drugs for nonclinical and clinical investigation in early stages are developed based on these knowledge.

The current realization demonstrated that mesenchymal characteristics of TNBC cells with paclitaxel-resistantance, and these cells has higher mesenchymal markers expression, including N-cadherin and vimentin. Thus, through inducing inhibition of TGF-β signaling pathway can promote chemotherapeutic effect on anti-tumor and reduce metastasis after paclitaxel therapy with kinase inhibitor for TGF-β1 receptor [27]. Therefore, it indicates that the novel potential strategy for TNBC therapy through combination with kinase inhibitors of TGF-β and its receptor.

Many data confirmed that microRNAs have crucial effect on activity of epithelial or EMT [28, 29]. So, in our investigation, the relationship between down-expression of miR-425 and TNBC was identified by comparing miRNA expression profiles from breast cancer tissues of patients with TNBC. Over-expression of miR-425 played an important regulator in negative feedback for TGF-β signaling and canonical pathway through suppressing expression of TGF-β1 and its changes associated with EMT in development of TNBC.

The system of tumor vessel includes pericytes, smooth muscle cells and, epithelial cells (ECs), which plays an important role in tumors progression and angiogenesis activation, such as breast cancer. The EMT process of ECs of breast cancer is conduced to the progression of cancer, that has been identified by recent researches. Therefore, the human breast cancer cell lines MDA-MB-231 and Hs 578T of ECs were choosed to address these process.

## Materials and Methods

### Specimens of patients

The tissues of patients with TNBC were obtained from January 2014 to December 2017. All these patients were performed with mastectomy for BC, and did not receive any treatment before hospital. At the operation day, samples of tumor tissues, para-tumor tissues and non-tumor tissues were collected and handled immediately, respectively. The para-tumor is adjacent tissue of tumor.

This investigation and the utilization of human specimens in this work had been approved by the Ethics Committee of our hospitals (Beijing Obstetrics and Gynecology Hospital, Capital Medical University, Beijing, China) on the basis of the Declaration of Helsinki. We clearly confirmed that the informed consents were obtained from all patients. Moreover, the recorded and documented of participant consents were kept in our hospital.

### Extraction of miRNAs and next-genersequencing

The miRNA-sequencing test was conducted after extraction of RNAs with TRIzol method and miRNeasy Mini Kit (Cat. No. 74104; Invitrogen·Thermo Fisher Scientific, Inc., Waltham, MA, USA). Furthermore, total RNA was harvested from tissues for RT-qPCR assay with mirVana™ PARIS™ Kit (Cat. No. AM1556; Invitrogen·Thermo Fisher Scientific, Inc, Waltham, MA, USA). Hybridization and ligation were conducted using Adaptor Mix. Moreover, microRNA sequencing RNAs using Illumina HiSeq 2000 platform was conducted after reversely transcribe of these RNAs.

### Data extraction and analysis

All the mature miRNAs sequences were identified through library preparation of reference sequences with miRBase (version 19.0) (http://www.mirbase.org/) [30]. Given that the length of minimum read was 50 nucleotides, specific adapters were used to anneal to their 3’ ends and extended the entire small RNAs during the library preparation. Thus, we removed the adapters and obtained the potential miRNAs with the length of 15-30 nucleotides for further analysis using cut adapt software in silico [31]. Subsequently, the sequences were perfectly matched with the prepared reference library with Bowtie (version 0.12.7) (https://sourceforge.net/projects/bowtie-bio/files/bowtie/0.12.7/) [32]. Then the numbers of mapped reads were counted for every miRNA. Acquired data of all samples were normalized with RPM (Reads per Million) [33].

### RNA Extraction

TRIzol method was used to extract total RNA from tissues based on the operation manual. Then, smaller RNA (<200 nt) was removed with mirVana RNA isolation kit (Cat. No. AM27828; Invitrogen·Thermo Fisher Scientific, Inc., Waltham, MA, USA) according to the instructions for manufacturer.

### Culture of Cell

The cell lines of human TNBC, including MDA-MB-231 (ATCC^®^ HTB-26^TM^) and Hs 578T (ATCC^®^ HTB-126^TM^) were purchased from ATCC (American Type Culture Collection), and then cultured in DMEM with 10% FBS accompanied by Streptomycin and Penicillin (bi-antibiotics) with atmosphere (5% CO_2_) at 37 °C.

### siRNA for TGF-β1

The well known siRNA of TGF-β1 supported by references [26] had been used in our study, which had been used as a positive in our study. The siRNA duplexes for TGF-β1 sequences are 5’-GAAGCGGACUACUAUGCUAUU-3’ (upstream) and 5’-UUCUUCGCCUGAUGAUACGAU-3’ (downstream).

### qRT-PCR

The miR-425 expression level was assessed with TaqMan miRNA assays in tissues. The data analysis was performed with 2^−ΔΔ^Cq method (Livak 2011). The qRT-PCR assay was conducted using SYBR Green kit (Cat. No. QR0100-1KT; Sigma-Aldrich; Merck KGaA, darmstadt, Germany). The thermocycling conditions for qPcR were as follows: Pre-denaturation at 95°C for 30 sec, followed by 40 cycles of denaturation at 95°C for 5 sec and extension at 60°C for 30 sec.

The primer sequences of miR-425 were 5′-ACACTCCAGCTGGGAATGACACGATCACTCC-3′ (upstream) and 5′-TGGTGTCGTGGAGTCG-3′ (downstream), respectively [34]. Moreover, the primer sequences of U6 were 5′-TGACCTGAAACATACACGGGA-3′ (upstream) and 5′-TATCGTTGTACTCCACTCCTTGAC-3′ (downstream), respectively.

The primer sequences of TGF-β1 were 5′-CAACACGATGCTTGAAGGTAACG-3′ (upstream) and 5′-TCCAGAGAGATGATTGCCGAGG-3′ (downstream), respectively. The primer sequences of VE-cadherin were 5′-GGATGCAGAGGCTCACAGAG-3′ (upstream) and 5′-CTGGCGGTTCACGTTGGACT-3′ (downstream), respectively. The primer sequences of E-cadherin were 5′-TACACTGCCCAGGAGCCAGA-3′ (upstream) and 5′-TGGCACCAGTGTCCGGATTA-3′ (downstream), respectively. The primer sequences of N-cadherin were 5′-GGACAGTTCCTGAGGGATCA-3′ (upstream) and 5′-GGATTGCCTTCCATGTCTGT-3′ (downstream), respectively. The primer sequences of Vimentin were 5′-GGCTCAGATTCAGGAACAGC-3′ (upstream) and 5′-GCTTCAACGGCAAAGTTCTC-3′ (downstream), respectively. The primer sequences of α-SMA were 5′-TGTTCCAGCCGTCCTTCAT-3′ (upstream) and 5′-GGCGTAGTCTTTCCTGATG-3′ (downstream), respectively. The primer sequences of SMAD3 were 5′-AGCACACAATAACTTGGACC-3′ (upstream) and 5′-TAAGACACACTGGAACAGCGGATG-3′ (downstream), respectively.

Moreover, the primer sequences of GAPDH (glyceraldehyde-3-phosphate dehydrogenase) were 5′-CTCATGACCACAGTCCATGCC-3′ (upstream) and 5′-GGCATGGACTGTGGTCATGAG-3′ (downstream), respectively.

### Reporter gene assays

The online software Targetscan (version 7.2) (http://www.targetscan.org/vert_72/) was firstly used to predict the microRNA which could regulate the expression of TGF-β1. Therefore, miR-425 was acquired from the predict result. Furthermore, the 3′UTR mRNAs sequence of TGF-β1 was cloned into pmirGLO vector with luciferase reporter firefly and renilla. The control was used with reverse orientation of TGF-β1 3′UTRs mRNA. The complementary region sequence in TGF-β1 3′UTR was GGACUGCGGAUCUCU**GUGUCAU**U (**Figure 1**), which located at position 151-157, and corresponding miR-425 seed sequence AGUUGCCCUCACUAG**CACAGUA**A. Site-Mutation kit (Cat. no. E0554S; New England Biolabs Ltd, UK) was used to constructed TGF-β1 3′UTR mutated vector. Finally, Dual-Glo ^®^ Luciferase Assay System (Cat. no. E2940; Promega corporation, Madison, WI, USA) was utilized to analyze the activity of luciferase with analyzer VICTOR.

**Figure1.**
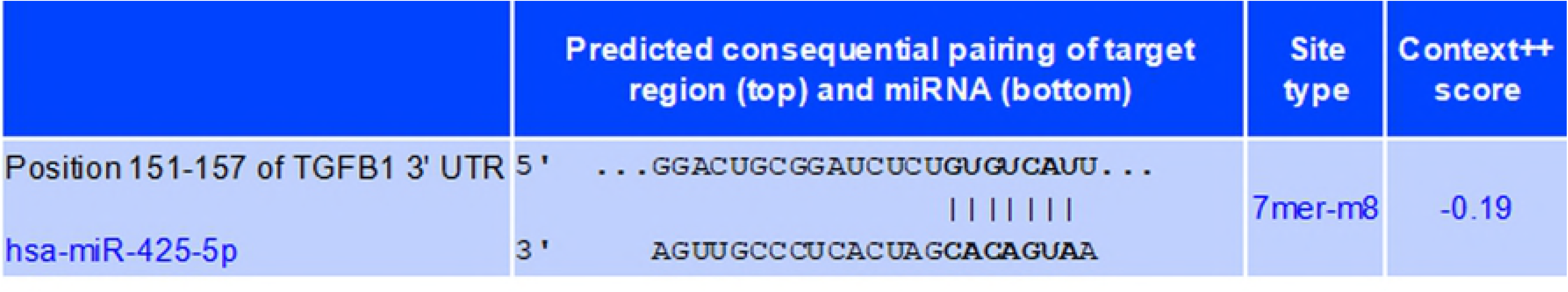
Predicted consequential pairing of target region. The online software Targetscan (version 7.2) was firstly used to predict the microRNA which could regulate the expression of TGF-β1. Therefore, miR-425 was acquired from the predict result. The complementary region sequence in TGF-β 1 3′ UTR was GGACUGCGGAUCUCUGUGUCAUU, which located at position 151-157, and corresponding miR-425 seed sequence AGUUGCCCUCACUAGCACAGUAA.

### The mimics of miR-425 and its inhibitors

The mimics of miR-425 and its inhibitors were transfected with Lipofectamine 3000. Moreover, the mimics sequences of miR-425 were 5′-AAUGACACGAUCACUCCCGUUGA-3′ [34]. Moreover, the control sequence of miR-425 mimics were 5’-UUCUCCGAACGUGUCACGUTT-3’. The sequence of miR-425 agomir was 5′-a_S_a_S_ugacacgaucacucccgu_S_u_S_g_S_a_s_-3′. The sequence of NC (negative control) was 5′-u_S_u_S_cuccgaacgugucacg_S_u_S_t_S_t_S_-3′, while it was not consistent with sequences of human genome.

### Western blot

A lysis buffer (1% NP-40, 0.1% SDS, 150 mM NaCl, 0.02% NaN_3_, and 50 mM Tris pH 8.0) containing cocktail of inhibitor for protease and phenylmethylsulfonyl fluoride (1 mM) was used to lysate cells. Western blotting assays were performed with Biolad Protean II minigel system. Then, 50 μg protein sample was injected in every gel (12%) well, and then shifted to PVDF membrane after electrophoresised. 1× TBST (5% dry skimmed milk) was utilized to dispute the non-specific binding. Then these membranes were sequentially dealt with primary and secondary antibody, respectively. The membranes were incubated with primary antibodies at 4° C overnight. The membranes were incubated with secondary antibodies with secondary antibodies goat anti-rabbit immunoglobulin G (IgG)-horseradish peroxidase (HRP) (sc-2004; 1:5,000 dilution; Santa Cruz Biotechnology, Inc.) or goat anti-mouse IgG-HRP (sc-2005; 1:5,000 dilution; Santa Cruz Biotechnology, Inc.) for 1 h at room temperature. Furthermore, these membranes were determined with enhanced ECL (chemiluminescence) reagent of Hyperfilm ECL kit (Cat. no. 28906835; Amersha·GE Healthcare, chicago, IL, USA), and exposed using x-ray film.

The primary antibodies included TGF-β1 (Santa Cruz, CA, USA; sc-130348, 1: 1200), α-SMA (CST 19245, dilution 1 : 1200) and SMAD 3 (C67H9) (CST 9523, dilution 1 : 1500), p-SMAD 3 (CST-9520, dilution 1 : 1000), VE-cadherin (CST2158, dilution 1 : 1000), E-cadherin (24E19) (CST 3195, dilution 1 : 800), N-cadherin (D4R1H) (CST 13116, dilution 1 : 1200), Vimentin (D21H3) (CST 5741, dilution 1 : 1000), β-actin (13E5) (CST 4970S, dilution 1 : 4000) (Cell Signaling, Danvers, MA, USA).

### Analysis of cell proliferation and migration

The proliferation ability of cells was assessed using MTS kit (Cat. no. G3580; Promega corporation, Madison, WI, USA). Subculture the stock cultures of cells to 2×10^4^ cells/ml in medium containing 50µM 2-mercaptoethanol (2-ME), when a density of 2×10^5^ cells/ml is reached, add mimics, inhibitors of miR-425 or its control, respectively. Wash the cells twice with the above medium centrifugation at 300×g for 5 minutes. 4. Determine cell number and viability (by trypan blue exclusion), and resuspend the cells to a final concentration of 1×10^5^ cells/ml in the above medium. Dispense 50µl of the cell suspension (5,000 cells) into all wells of the 96-well plate prepared. The total volume in each well should be 100µl. Incubate the plate at 37°C for 48 h in a humidified, 5% CO_2_ atmosphere. Add 20µl per well of CellTiter 96^®^ AQueous One Solution Reagent. Incubate the plate at 37° C for 2 h in a humidified, 5% CO_2_ atmosphere. Record the absorbance at 490nm using a 96-well plate reader.

The ability of cells migration was detected using wound scratch assay using CytoSelect™ 24-Well Wound Healing Assay kit (Cat. no. CBA-120-5; Cell Biolabs, Inc., San Diego, CA, USA). Allow the 24-well plate with CytoSelect™ Wound Healing Inserts to warm up at room temperature for 10 minutes. Using sterile forceps, orient the desired number of inserts in the plate wells with their “wound field” aligned in the same direction. Ensure that the inserts have firm contact with the bottom of the plate well. Create a cell suspension containing 1×10^6^ cells/ml in media containing 10% fetal bovine serum (FBS). Add 500 µL of cell suspension to each well by carefully inserting the pipet tip through the open end at the top of the insert. For optimal cell dispersion, add 250 µL of cell suspension to either side of the open ends at the top of the insert. Take care to avoid bumping and moving the inserts. Incubate cells in a cell culture incubator overnight or until a monolayer forms. Carefully remove the insert from the well to begin the wound healing assay. Use sterile forceps to grab and lift the insert slowly from the plate well. Avoid twisting the insert as this will damage the wound field. Slowly aspirate and discard the media from the wells. Wash wells with media to remove dead cells and debris. Finally, add media to wells to keep cells hydrated. Visualize wells under a light microscope. Repeat wash if wells still have debris or unattached cells. When washing is complete, add media with FBS and/or compounds to continue cell culture and wound healing process. Incubate cells in cell culture incubator. For best results, use a reticle with micrometer measurements to create a defined surface area in order to monitor the closing, or “healing” of the wound. Focus on the center of the wound field. Create the defined surface area by multiplying the width of the wound field (0.9 mm) by the length. Monitor the wound closure with a light microscope or imaging software. Measure the percent closure or the migration rate of the cells into the wound field. Wound healing results can be visualized with phase contrast.

Ten fields per experimental condition were randomly selected and micrographed with IX71 microscope (Olympus, japan) to quantify cells of migration. Images represent at least three independent experiments.

### Matrigel invasion assay

Matrigel was dissolved overnight at 4°C and then diluted with serum-free DMEM (1:3). The diluted Matrigel was added to the upper chamber of the Transwell to form the membrane in three additions (15, 7.5 and 7.5 μl), with 10 min. between each addition. The Matrigel was evenly spread and covered all micropores at the bottom of the upper chamber. The two cell lines were collected at 48 h after transfection to prepare cell suspensions. Cell suspensions were seeded into the upper well, and 0.5 ml of DMEM containing 10% FBS was added to the lower chambers (24-well plates). The cells were cultured routinely for 48 h at 37°C with 5% CO_2_. After using a cotton swab to gently wipe off the remaining free cells in the upper well, the well membrane was collected and fixed for 15 ~ 20 min. in 95% ethanol. After rinsing with clean water, the membrane was stained for 10 min. and washed again with clean water. The membrane was then observed under a high-powered inverted microscope to count the number of cells on the membrane. The invasion ability of the cells was determined by counting the number of cells that passed through the Matrigel, and the average number from five high-magnification fields of view was used for each sample.

### Animals

Nude mice (purchased from Laboratory Animal Sciences, Capital Medical University, Beijing, China) were used to conduct our investigation at this work. A total of 12 nude mice,female, were randomly divided into 2 groups, and the number of males and females in each group was guaranteed to be equal. The experiment was initiated with 6 week-old mice weighing 20-25 g. They were induced to generate xenograft model with injection of MDA-MB-231 cells, and then housed of Laboratory Animal Center, Capital Medical University, Beijing, China. Mice were maintained at 20-24 ºC (temperature controlled) with 12 h light/dark cycle and standard laboratory diet in a pathogen-free environment. All experiments related to animals were performed based on Laboratory Animal Center, Capital Medical University, Beijing, China, Ethical Committee Acts. The experimental mice were dealt with according to the standards supported by the Animal Protection Committee of Capital Medical University. Moreover, the ethical approval was obtained for the animal experiments conducted in your study from Animal Protection Committee of Capital Medical University.

### Experiment in vivo

The experiment was initiated with 6 week-old of mice, which weight was 20-25 g. Then, 1 ml suspension of MDA-MB-231 cells (1×10^6^ cells/ml) was injected into subcutaneous tissue of nude mice at right flanks (n=6/group). While tumor size of mice reached 3-8 mm, the experiment initiated as the 1st day. The agomir of miR-425 were formulated (1.0 mg/ml in PBS). The nude mice were categorized into experiment and control groups at random, then injected with agomir of miR-425 (5 µl) or it control once every three days for 6 times and terminated on the 18th day. Then, these mice were sacrificed at the 21th day, and tumor mass was dissected from each mouse. These tumors were cut into two sections, then the volume of tumors were determined every 3 days base on the formulate width^2^ × length × 0.5. Moreover, the expression of TGF-β1 were determined with western blotting and immunohistochemical staining assay, respectively.

The dose of urethane used for euthanasia of mice is (at 1 g/kg), the following methods were used to evaluate consciousness or confirm death. We verified animal death in our study after the administration of urethane by observing lack of a heartbeat, lack of respiration, and lack of corneal reflex for more than 16 seconds. The signs of pain or disease, such as the earliest indicator in an animal of pain, distress, suffering, or impending death were considered humane endpoints in our study.

### Immunohistochemistry assay

Tissue samples were prepared using 10% neutral formalin, then embedded using paraffin after dehydration. Furthermore, it was cut into thick sections with 5 µM (Marcela et al. 2018), dewaxed and hydrated. After antigen retrieval, incubation with primary antibody for these sections was performed. The sections were incubated with primary antibodies TGF-β1 (Santa Cruz, CA, USA; sc-130348, 1: 200), or SMAD 3 (C67H9) (CST 9523, dilution 1 : 200) at 4°C overnight, respectively. The sections were incubated with a DAB substrate kit (Cat. no. 34002; Thermo Scientific™ Pierce™, San Jose, CA, USA) after incubation with the secondary antibody. The membranes were incubated with secondary antibodies with secondary antibodies goat anti-rabbit immunoglobulin G (IgG)-horseradish peroxidase (HRP) (sc-2004; 1:1,000 dilution; Santa Cruz Biotechnology, Inc.) or goat anti-mouse IgG-HRP, (sc-2005; 1:1,000 dilution; Santa Cruz Biotechnology, Inc.) for 1 h at room temperature. These sections were assessed, and then photographed with microscope at 200× (IX71; Olympus, Japan). Per sample was token for 3-5 shots. The software Image-Proplus (version 6.0) was used measure proteins expression through detecting optical density in each photograph.

### Statistics

Welch *t*-test-paired was performed to compare isomiRs deregulation and selection of miRNAs between samples of patients and control. The errors were evaluated with false discovery rate (FDR) due to multiple comparisons. On the basis of the expression profiles, Hierarchical clustering of selected miRNAs was conducted with Ward’s agglomeration. Target Rank software (http://genes.mit.edu/targetrank/) was performed to identify the significantly deregulated each seed sequence of miRNA in target genes between samples of patients and control [35].

All data with normally distributed was represented as mean ± SD (standard deviation). The differences between two groups were analyzed with Student’s *t*-test. Kruskal-Wallis ANOVA method was used to analyze abnormally distributed data among groups. When the variances were equal, the least signidicant difference post hoc test was used, whereas when the variances were not uniform, Dunnett's post hoc test was used. SPSS (version 18.0; SPSS, Inc., chicago, IL, USA) soft was performed for entire statistical analyses. While *P*< 0.05 indicated statistically significant.

## Results

### Differential miRNA expression profiling

To assess the differential expression of miRNA profiles between tumorous and para-tumorous tissues, miRNA-sequencing experiments were conducted on the total RNA obtained from samples. Among the results of miRNAs-sequencing, 12 were differentially expressed between tumorous and para-tumorous tissues. Compared with para-tumorous tissues, 9 miRNAs were upregulated and 3 miRNAs were down-regulated in patients with TNBC (**Table 1**).

**Table 1.** Differentially expressed **miRNAs**m tumorous and tumorous tissue., (fold-changc>2.0 and P-value<0.01).

| miRNA | Fold-change (tumorous/para-tumorous tissues) | *t* value | *P* value |
| --- | --- | --- | --- |
| miR-148b | 5.73 | 8.381 | 7.72x10<sup>-4</sup> |
| miR-34a | 4.08 | 6.935 | 9.33x10<sup>-3</sup> |
| miR-196a | 4.01 | 7.220 | 7.00x10<sup>-3</sup> |
| miR-205 | 3.91 | 9.272 | 5.46x10<sup>-3</sup> |
| miR-328 | 3.18 | 2.702 | 6.96x10<sup>-3</sup> |
| miR-122 | 2.64 | 7.254 | 1.77x10<sup>-3</sup> |
| miR-155 | 2.45 | 3.823 | 5.44x10<sup>-3</sup> |
| miR-202 | 2.36 | 6.256 | 7.68x10<sup>-3</sup> |
| miR-let-7a | 2.34 | 3.293 | 4.42x10<sup>-3</sup> |
| miR-7 | 0.35 | -0.391 | 5.49x10<sup>-3</sup> |
| **miR-425** | 0.18 | -0.468 | 7.53x10<sup>-3</sup> |
| miR-31 | 0.14 | -0.326 | 2.07x10<sup>-3</sup> |

### The expression level of miR-425 in non-tumorous, para-tumorous, and tumorous tissues

Furthermore, the relative miR-425 expression levels in non-tumorous, para-tumorous, and tumor tissues (n=60) were determined using qRT-PCR method. Our work identified the relative miR-425 expression level was obviously less than that of non-tumorous or para-tumorous tissues. However, there was no obvious difference between para-tumorous and non-tumorous tissues (**Figure 2**).

**Figure2.**
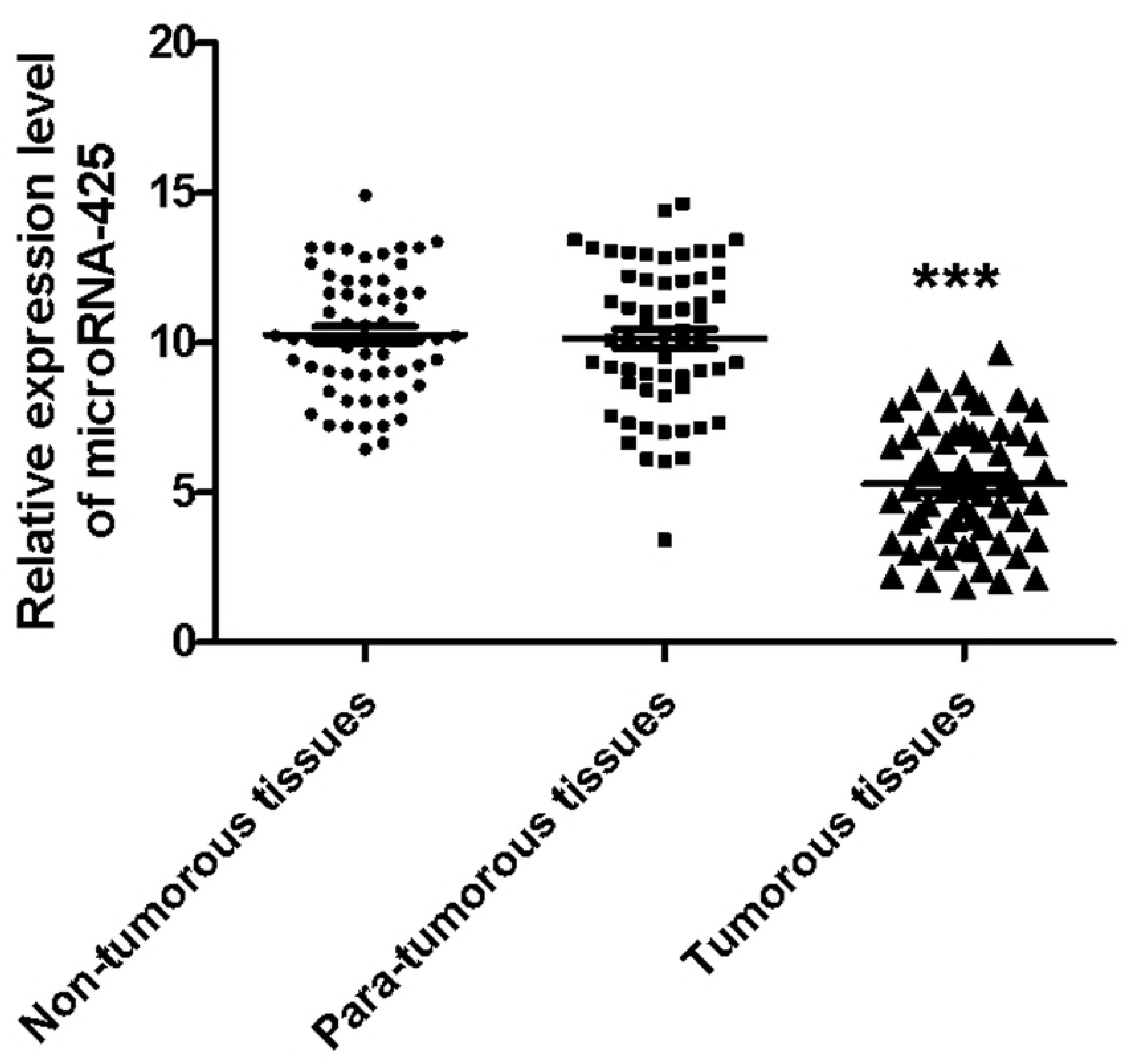
The expression level of miR-425 in non-tumor, para-tumor, and tumor tissues. The relative miR-425 expression levels in non-tumor, para-tumor, and tumor tissues (n=60) were determined using qRT-PCR method. This result identified the relative miR-425 expression level in tumor tissues was obviously less than that of non-tumor or para-tumor tissues (*P*< 0.001). However, there was no obvious difference between para-tumor and non-tumor tissues. \*\*\**P*<0.001.

### Expression levels of miR-425 were determined in cell lines of human TNBC

Moreover, the relative miR-425 expression levels were investigated in two cell lines of human TNBC. Compared with basal epithelial phenotype cell line Hs 578T, the relative miR-425 expression levels were significantly higher in mesenchymal phenotypic cell line MDA-MB-231 (*P*< 0.001) (**Figure 3**).

**Figure3.**
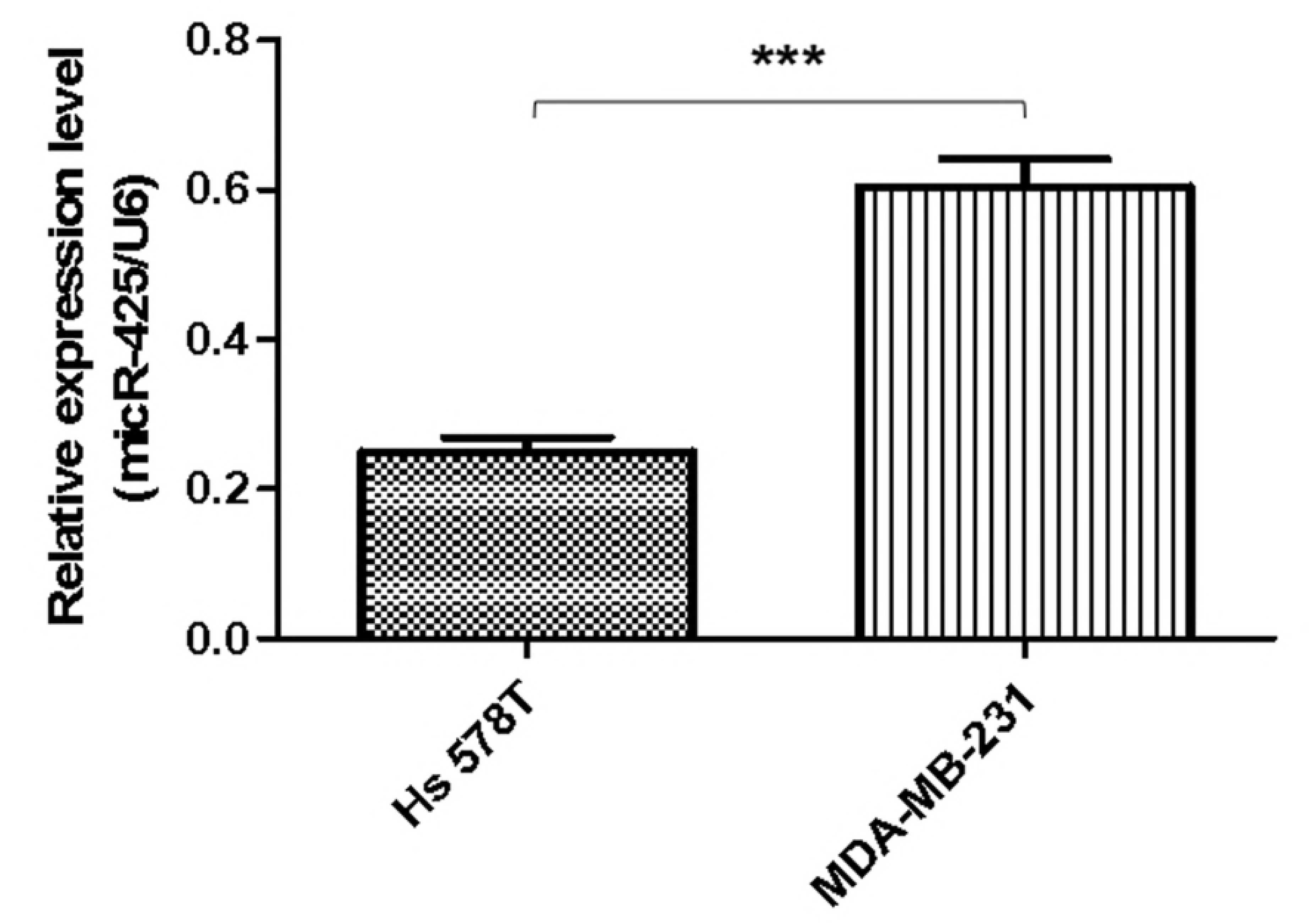
Expression levels of miR-425 were determined in cell lines of human TNBC. The relative miR-425 expression levels were investigated in six cell lines of human TNBC. Compared with basal epithelial phenotype cell line Hs 578T, the relative miR-425 expression levels were significantly higher in mesenchymal phenotypic cell line MDA-MB-231 (*P*< 0.001). The data are presented as means ± SD from at least three independent experiments. \*\*\**P*<0.001.

### MiR-425 inhibited TGF-β1/SMAD3 pathway and mesenchymal markers expression in TNBC cell lines

At first, the target of miR-425 were predicted with Targetscan, an online software. Therefore, miR-425 was found that it is the possibly target of 3’ UTRs untranslated region of TGF-β1 mRNA (**Table 1**). Further experiments were conducted using luciferase reporter gene system to identify whether miR-425 could straight bind with 3’ UTRs of TGF-β1 mRNA. The data indicated 3’ UTR luciferase activities of TGF-β1 significantly down-regulated in MDA-MB-231 cell line dealt with miR-425 (**Figure 4A**). However, it did not show obviously change in the group of TGF-β1 mutation (**Figure 4B**). Moreover, miR-425 did not bind straight to 3’ UTRs of E-cadherin (**Figure 4C**), VE-cadherin (**Figure 4D**), N-cadherin (**Figure 4E**), Vimentin (**Figure 4F**), α-SMA (**Figure 4G**), and SMAD3 (**Figure 4H**), respectively. These data identified that miR-425 targeted to TGF-β1 mRNAs. The well known siRNA of TGF-β1 supported by references [26] had been used in our study, which had been used as a positive in our study. It could inhibit the TGF-β 1/SMAD3 signaling pathway in TNBC cell lines (**Figure 5**).

**Figure4.**
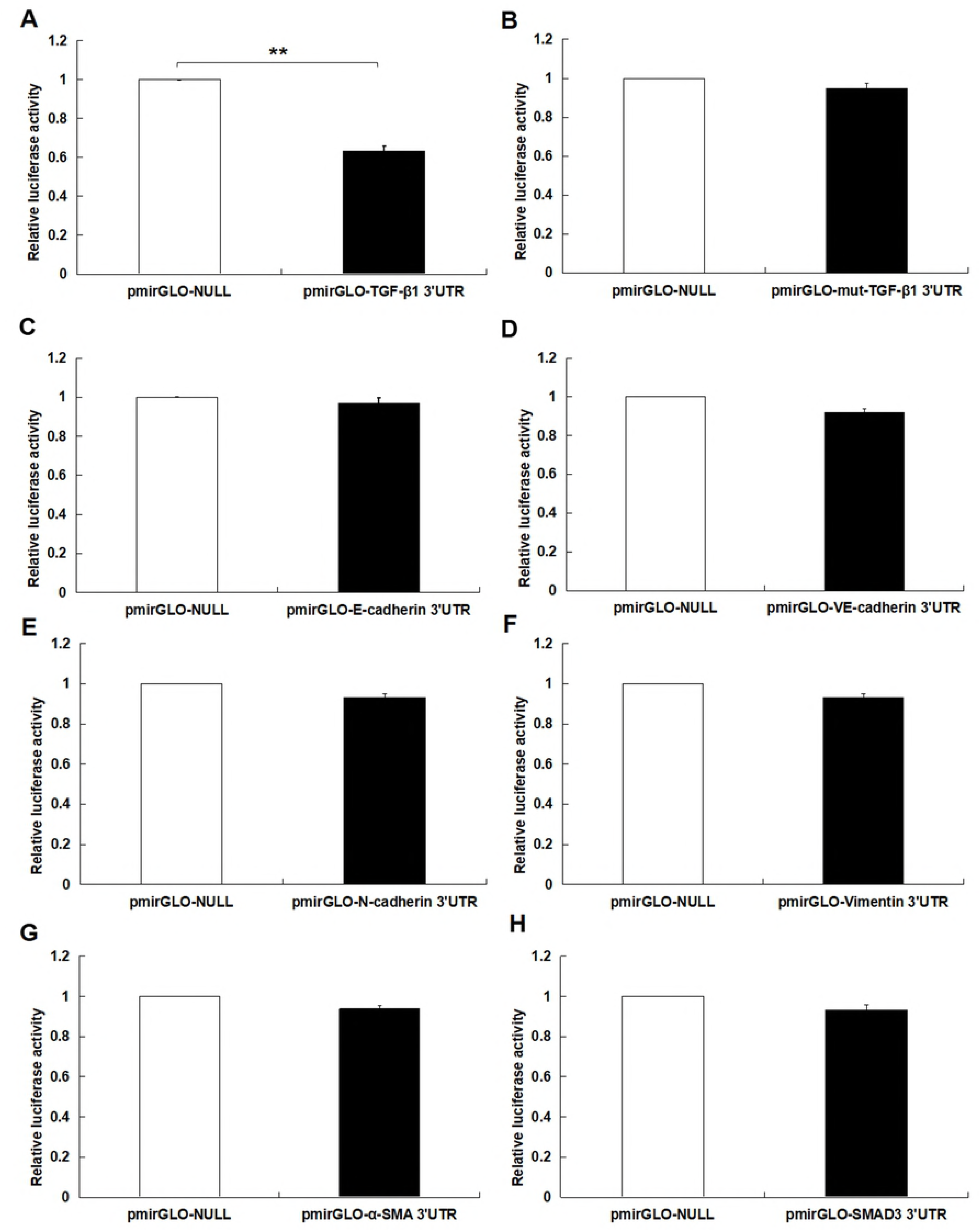
MiR-425 inhibited TGF-β1 expression in TNBC cell lines. The target of miR-425 were predicted with Targetscan, the online software. Therefore, miR-425 was found that possibly aimed to 3’ UTRs untranslated region of TGF-β1 mRNA. Further experiments were conducted using luciferase reporter gene system to identify whether miR-425 could straight bind with 3’ UTRs of TGF-β1 mRNA. The data indicated 3’ UTR luciferase activities of TGF-β1 significantly down-regulated in MDA-MB-231 cell line dealt with miR-425 (**A**). However, it did not show obviously change in the group of TGF-β1 mutation (**B**). Moreover, miR-425 did not bind straight to 3’ UTRs of E-cadherin (**C**), VE-cadherin (**D**), N-cadherin (**E**), Vimentin (**F**), α-SMA (**G**), and SMAD3 (**H**), respectively. These data identified that miR-425 targeted to TGF-β1 mRNAs. The data are presented as means ± SD from three independent experiments. \**P*<0.05, \*\**P*<0.01.

**Figure5.**
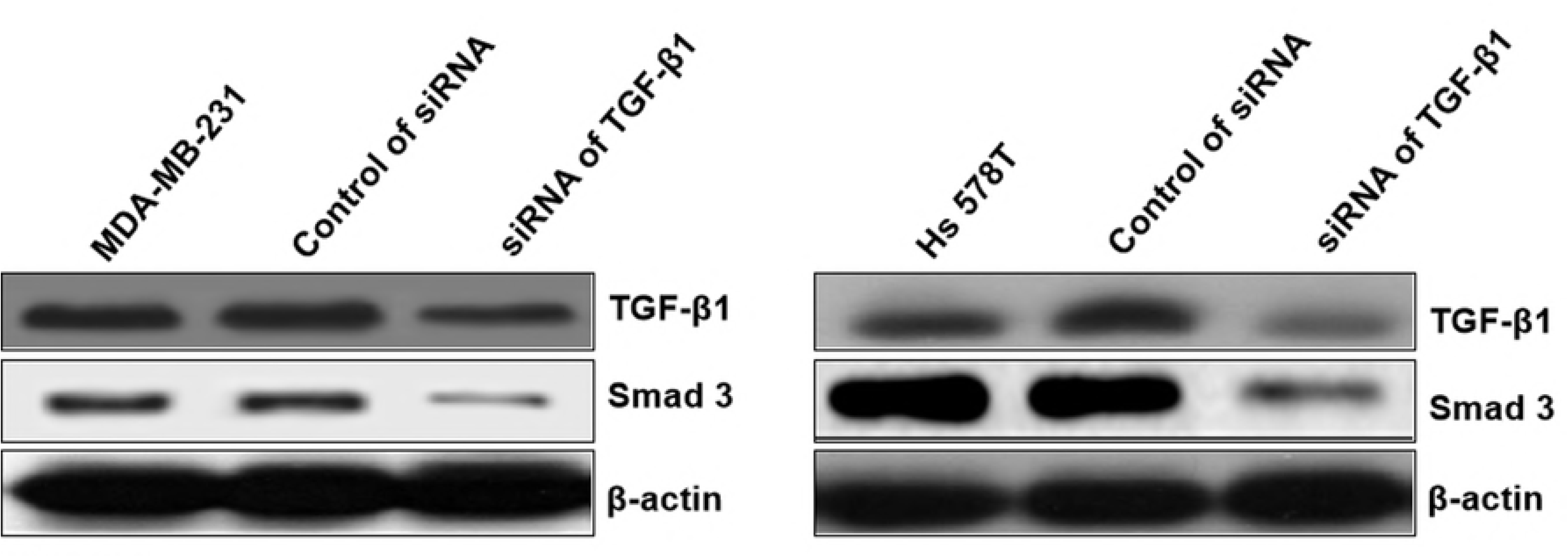
The siRNA of TGF-β1 inhibited TGF-β1/SMAD3 signaling pathway in TNBC cell lines. The well known siRNA of TGF-β1 supported by references [26] had been used in our study, which had been used as a positive in our study. It could inhibit the TGF-**β**1/SMAD3 signaling pathway.

To explore the roles of miR-425 on mRNAs and proteins expression of TGF-β1, α-SMA, Vimentin, N-cadherin, E-cadherin, VE-cadherin, and SMAD3 in human BC cell lines, the qRT-PCR assay and western blotting trial were performed. These results demonstrated that miR-425 restrained mRNAs and proteins expressions of TGF-β1 in MDA-MB-231 cell line. However, miR-425 could not suppress the mRNAs expression of α-SMA, Vimentin, N-cadherin, VE-cadherin, E-cadherin, and SMAD3 (**Figure 6A**). The western blotting result showed that miR-425 promoted proteins expression of E-cadherin and VE-cadherin, while suppressed proteins expression of α-SMA, Vimentin, N-cadherin, and SMAD3 (**Figure 6C**). Moreover, miR-425 inhibitors could promote expression of TGF-β1 mRNA in MDA-MB-231, but did not affect mRNAs expression of α-SMA, Vimentin, N-cadherin, VE-cadherin, E-cadherin, and SMAD3 (**Figure 6A**). The inhibitors of miR-425 could suppress the protein expression levels of N-cadherin, Vimentin, α-SMA, and SMAD3, but promote VE-cadherin and E-cadherin protein expression level (**Figure 6C**). In a word, the result indicated that, by binding with TGF-β1 3′-UTR, miR-425 could straight regulate its mRNA and protein expression, further affect protein expression of genes associated with EndMT process and TGF-β1/SMAD3 pathway. It was same as the results in cell line Hs 578T (**Figure 6B and 6D**).

**Figure6.**
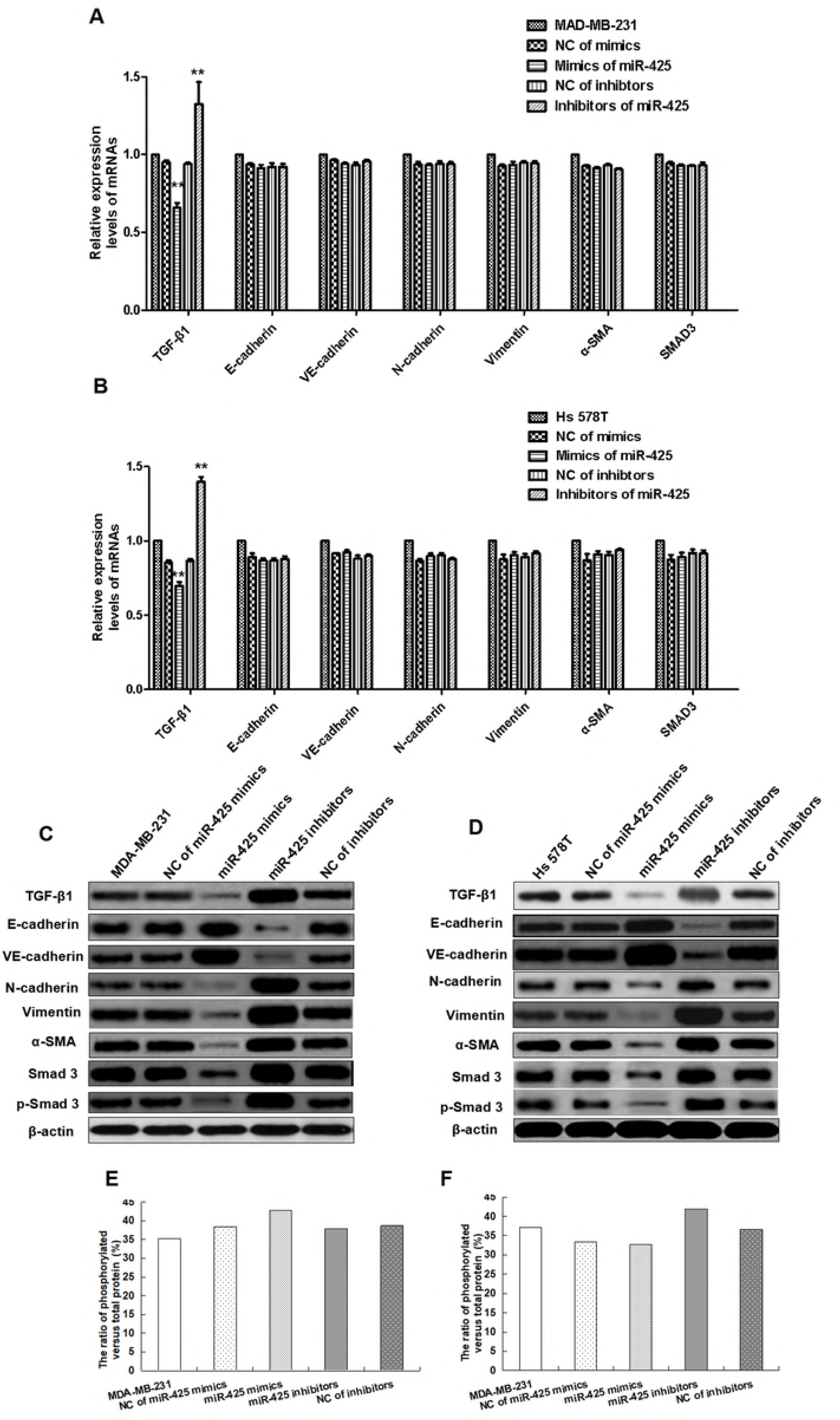
MiR-425 inhibited TGF-β1/SMAD3 pathway and mesenchymal markers expression in TNBC cell lines. To explore the roles of miR-425 on mRNAs and proteins expression of TGF-β1, E-cadherin, VE-cadherin, N-cadherin, Vimentin, α-SMA, total and phosphorylated SMAD3 (p-SMAD3) in human BC cell lines, the qRT-PCR assay and western blotting trial were performed. These results demonstrated miR-425 restrained TGF-β1 mRNAs and proteins expressions in MDA-MB-231 cell line. However, miR-425 could not suppress the expression of E-cadherin, VE-cadherin, N-cadherin, Vimentin, α-SMA, and SMAD3 mRNAs (**A**), but promote proteins expression of E-cadherin and VE-cadherin, while suppress proteins expression of N-cadherin, Vimentin, α-SMA, total and phosphorylated SMAD3 (p-SMAD3) (**C**). Moreover, miR-425 inhibitors could promote expression of TGF-β1 mRNA in MDA-MB-231, but did not affect mRNAs expression of E-cadherin, VE-cadherin, N-cadherin, Vimentin, α-SMA, total and phosphorylated SMAD3 (p-SMAD3) (**A**). The inhibitors of miR-425 could suppress the protein expression levels of N-cadherin, Vimentin, α-SMA, total and phosphorylated SMAD3 (p-SMAD3), but promote protein expression of E-cadherin and VE-cadherin (**C**). In a word, by binding with TGF-β1 3′-UTR, the result indicated miR-425 could straight regulate its mRNA and protein expression, further affect protein expression of genes associated with EndMT process and TGF-β1/SMAD3 pathway. It was same as the results in other cell line Hs 578T (**B and D**). The mesenchymal markers expression of N-cadherin, Vimentin, α-SMA, SMAD3 and p-SMAD3 in TNBC cell lines were inhibited significantly, while endothelial markers expression of E-cadherin and VE-cadherin were obviously promoted by mimics of miR-425 *in vitro* compared with control group (**C and D**). But, the expression of mesenchymal cell markers in TNBC cell lines were significantly promoted, while endothelial markers expression of E-cadherin and VE-cadherin were significantly suppressed by mimics of miR-425 in TNBC cell lines (**C and D**). The ratio of phosphorylated versus total protein did not show significant difference between each groups in MDA-MB-231 and Hs 578T cell lines (**E and F**). The data are presented as means ± SD from three independent experiments. \**P*<0.05, \*\**P*<0.01.

The mesenchymal markers expression of α-SMA, Vimentin, N-cadherin, SMAD3 and p-SMAD3 in TNBC cell lines were significantly inhibited, while endothelial markers expression of E-cadherin and VE-cadherin were obviously promoted by mimics of miR-425 *in vitro* compared with control group (**Figure 6C and 6D**). But, the mesenchymal cell markers expression of TNBC cells were significantly promoted, while endothelial markers expression of E-cadherin and VE-cadherin were significantly suppressed by inhibitors of miR-425 in TNBC cells (**Figure 6C and 6D**). The ratio of phosphorylated versus total protein did not show significant difference between each groups in MDA-MB-231 and Hs 578T cell lines (**Figure 6E and 6F**). Our results identified that miR-425 suppressed mesenchymal markers expression and enhanced endothelial markers expression in TNBC cell lines, especially for the mesenchymal phenotypic cell line MDA-MB-231. The expressions of mesenchymal markers N-cadherin, Vimentin, α-SMA, SMAD3 and p-SMAD3 in the mesenchymal phenotypic cell line MDA-MB-231 were higher than that in the basal phenotypic epithelial cell line Hs 578T, and all could be suppressed obviously by mimics of miR-425 in the two cell lines. Moreover, the expressions of endothelial markers E-cadherin and VE-cadherin in the mesenchymal phenotypic cell line MDA-MB-231 were less than that in the basal phenotypic epithelial cell line Hs 578T, and all two cell lines could be promoted significantly by mimics of miR-425.

### MiR-425 suppressed cell proliferation, migration, and invasion of TNBC cell lines

To assess the biofunctions of miR-425, over-expressed and down-expressed miR-425 was conducted, respectively. Furthermore, several molecular functional tests were performed in TNBC cells. The proliferation ability of TNBC cells was suppressed by miR-425 compared to that of control, and suppressed by inhibitors of miR-425 (**Figure 7A**). Moreover, the change of proliferation ability induced by mimics of miR-425 in mesenchymal phenotypic cell line MDA-MB-231 was significantly more than that of basal epithelial phenotype cell line Hs 578T, while the change between the two cell lines was statistically significant difference (*P*<0.05) (**Figure 7A**).

**Figure7.**
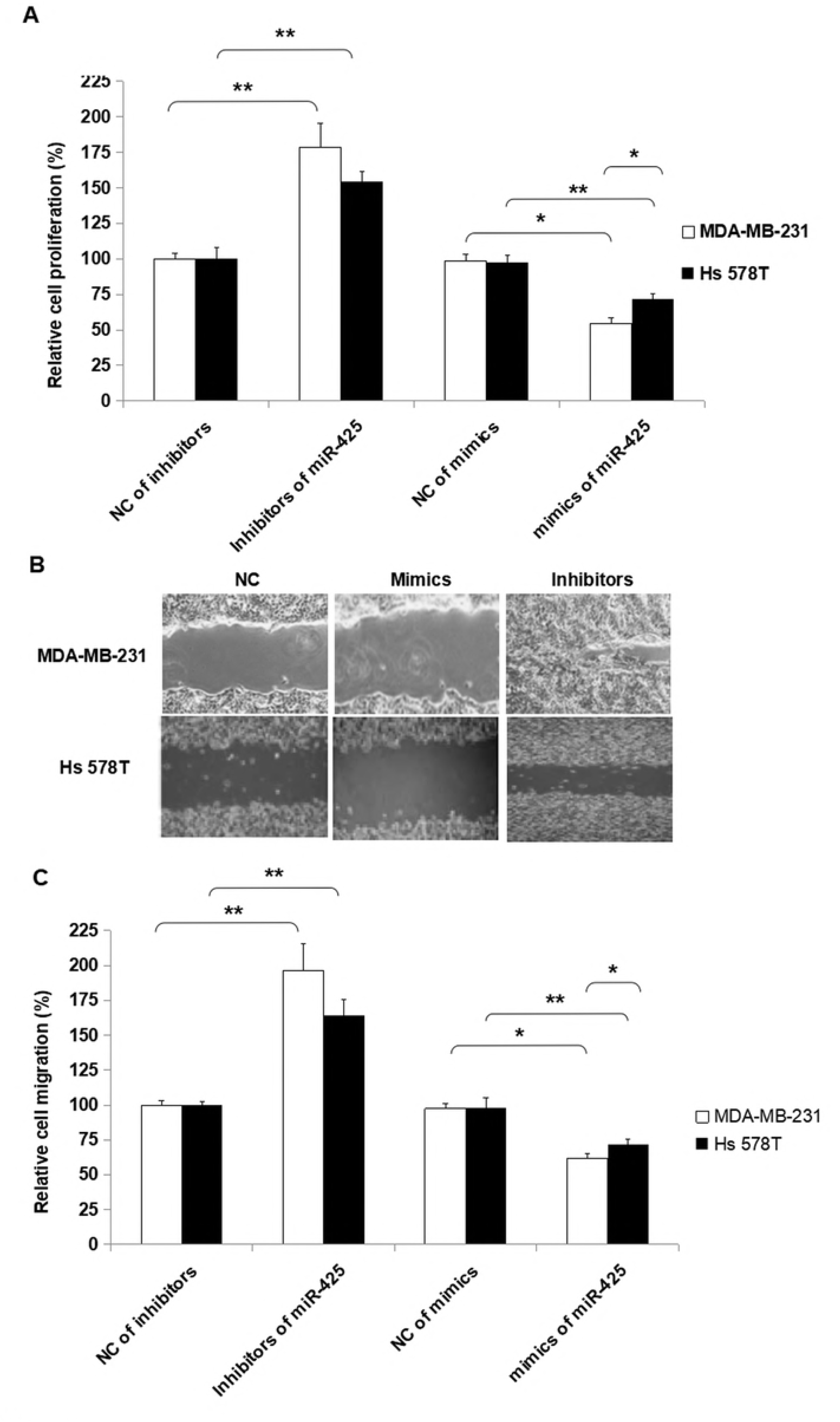
MiR-425 suppressed cell proliferation and migration of TNBC cell lines. To assess the biofunctions of miR-425, over-expressed and down-expressed miR-425 was conducted, respectively. Furthermore, several molecular functional tests were performed in TNBC cell lines. The proliferation ability of TNBC cell lines was suppressed by miR-425 compared to that of control. Moreover, the change of proliferation ability induced by mimics of miR-425 in mesenchymal phenotypic cell line MDA-MB-231 was significantly more than that of basal epithelial phenotype cell line Hs 578T, while the difference was statistically significant (*P*<0.05). Additionally, the inhibitors of miR-425 promoted proliferation ability of TNBC cell lines. Furthermore, the change of proliferation ability induced by inhibitors of miR-425 in mesenchymal phenotypic cell line MDA-MB-231 was higher than that of basal epithelial phenotype cell line Hs 578T, while the difference between the two type cell lines was not statistically significant (*P*>0.05). The data are presented as means ± SD from three independent experiments. \**P*<0.05, \*\**P*<0.01.

Additionally, the inhibitors of miR-425 promoted migration ability of TNBC cell lines, but the mimics of miR-425 suppressed the migration ability of TNBC cell lines (**Figure 7B and 7C**). Furthermore, the change of migration ability induced by inhibitors of miR-425 in mesenchymal phenotypic cell line MDA-MB-231 was higher than that of basal epithelial phenotype cell line Hs 578T, while the change between the two type cell lines was not statistically significant difference (*P*>0.05) (**Figure 7B and 7C**). It identified that miR-425 suppressed the process of proliferation and migration in TNBC cell lines, especially for mesenchymal phenotypic cell line MDA-MB-231. The hypothesis to explain these results was that the inhibition roles induced by miR-425 in proliferation and migration of TNBC cell line, mesenchymal phenotypic cell line MDA-MB-231 were much more than the basal epithelial phenotype cell line Hs 578T **(Figure 7)**. Moreover, it maybe the mesenchymal characteristics of MDA-MB-231 cell line was obviously more than Hs 578T cell line. Moreover, the inhibitors of miR-425 promoted invasion ability of TNBC cell lines, but the mimics of miR-425 suppressed the invasion ability of TNBC cell lines (Figure 8).

**Figure8.**
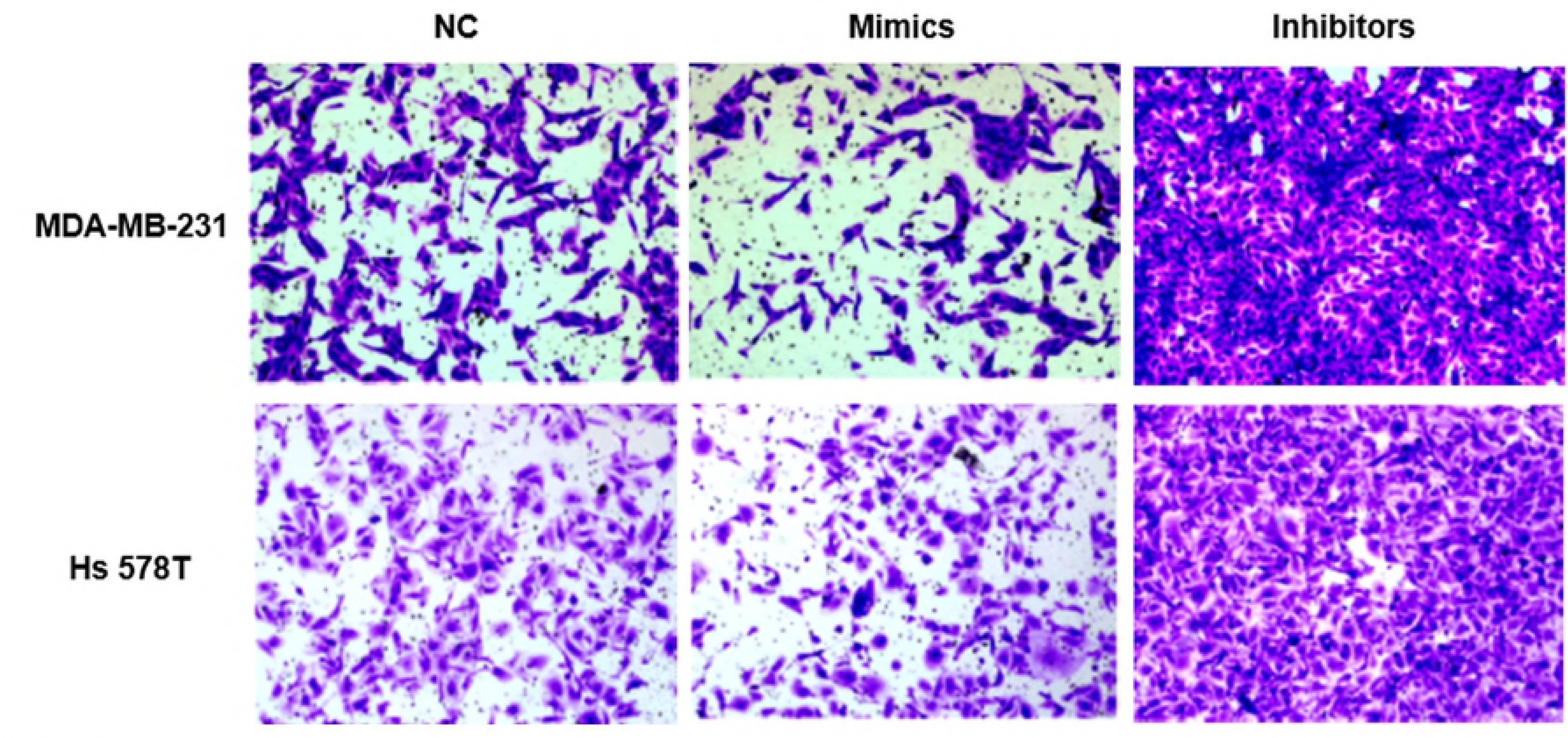
MiR-425 suppressed cell invasion of TNBC cell lines. Moreover, the inhibitors of miR-425 promoted invasion ability of TNBC cell lines, but the mimics of miR-425 suppressed the invasion ability of TNBC cell lines.

### MiR-425 agomir suppressed TNBC xenografts

To detect anti-tumor activity induced by miR-425 agomir *in vivo*, human TNBC xenografts were established with MDA-MB-231 cells. Our work demonstrated that the size of tumor (**Figure 9A**) and the average tumor weight (**Figure 9B**) of control mice was much larger than that of experiment mice treated with agomir of miR-425, and the difference between two groups had statistical significance. Moreover, the xenografts of control groups grew rapidly, but in the experiment mice treated with agomir of miR-425, the xenografts grew slowly *in vivo* (**Figure 9C**). The maximum diameter exhibited by a single subcutaneous tumor of control mice in our study was 1.23 cm.

**Figure9.**
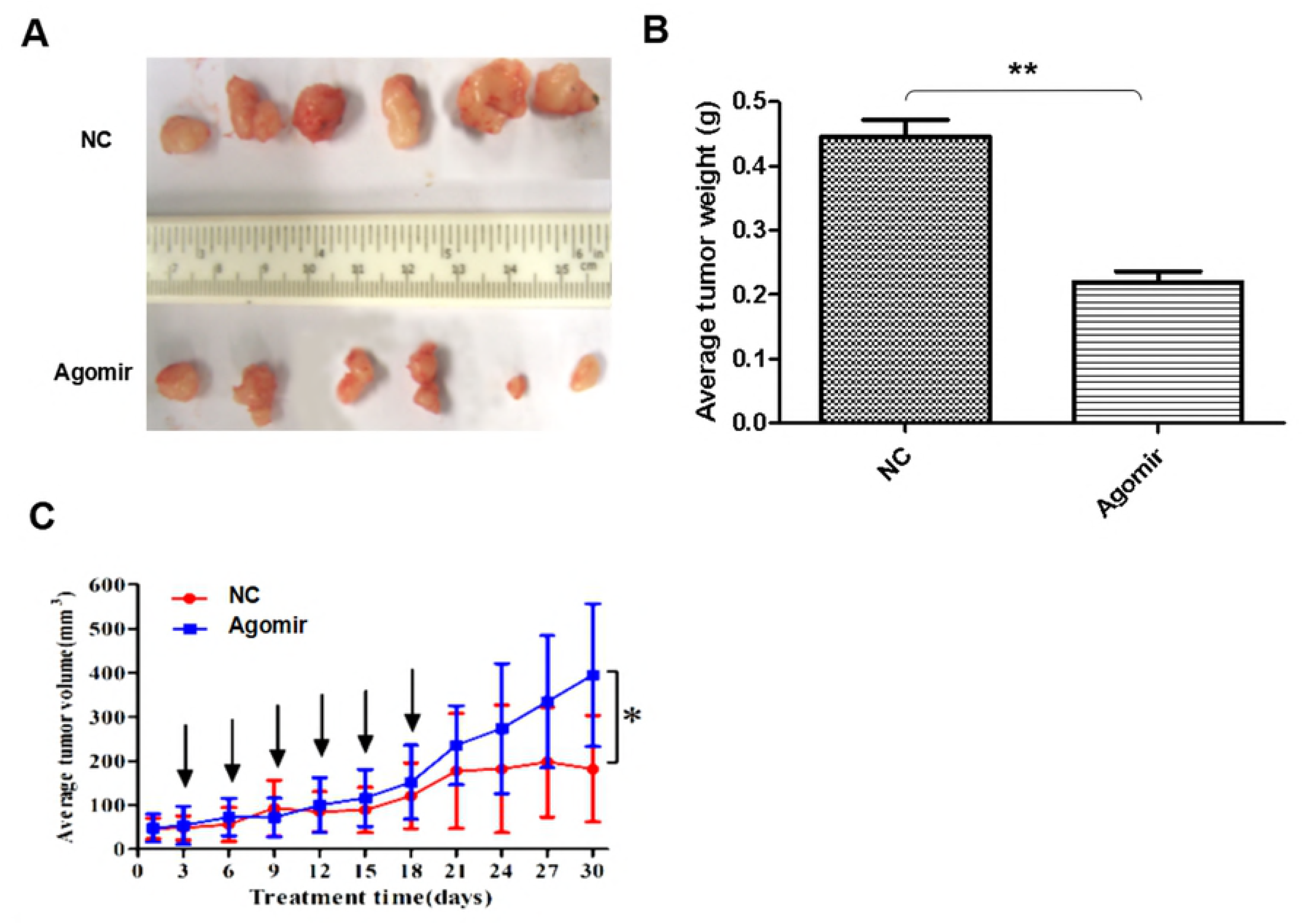
MiR-425 agomir suppressed TNBC xenografts. To detect anti-tumor activity induced by miR-425 agomir *in vivo*, human TNBC xenografts were established with MDA-MB-231 cells. Our work demonstrated that the size of tumor (**A**) and the average tumor weight (**B**) of control mice was much larger than that of experiment mice treated with agomir of miR-425, and the difference between two groups had statistical significance. Moreover, the xenografts of control groups grew rapidly, but in the experiment group, which was treated with agomir of miR-425, grew slowly *in vivo* (**C**). The data are presented as means ± SD from three independent experiments. \**P*<0.05, \*\**P*<0.01.

The expression proteins in xenografts were determined with immunohistochemical staining methods (**Figure 10A**) and western blotting assay (**Figure 10B)**, respectively. The results revealed that when compare with control group, treatment with agomir of miR-425 could down-regulate the expression of SMAD3 and TGF-β1 with a statistically significant difference, and suppressed mesenchymal markers expression of xenograft. The mesenchymal markers expression of α-SMA, Vimentin, N-cadherin, SMAD3, p-SMAD3 and endothelial markers E-cadherin, VE-cadherinin were determined in xenografts. The results indicated that compare with control group, treatment with agomir of miR-425 could obviously down-regulate the above mesenchymal markers expression in xenografts, while up-regulate expression of endothelial markers E-cadherin, VE-cadherinin. To sum up, our data confirmed that the agomir of miR-425 could inhibit development of TNBC associated with EMT through aiming TGF-β1/SMAD3 signaling pathway *in vivo*.

**Figure10.**
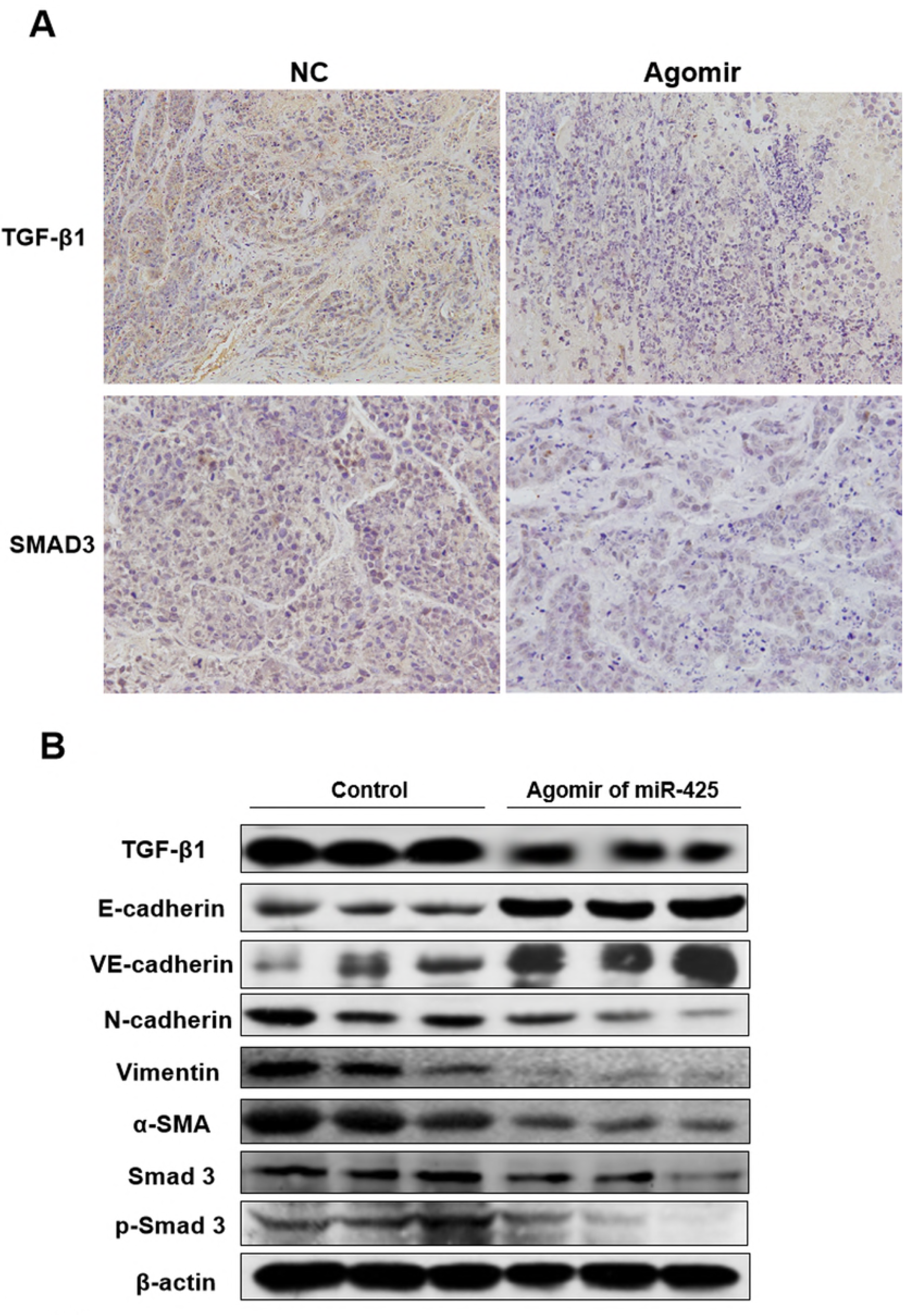
The miR-425 agomir suppressed expression of mesenchymal cell markers in xenografts. The expression proteins in xenografts were determined with immunohistochemical staining (**A**) and western blotting methods (**B)**, respectively. As we can see, the results revealed that when compare with control group (from left to right, control 1-3), treatment with agomir of miR-425 (from left to right, experiment 1-3) could down-regulate the expression of TGF-β1 and SMAD3 with a statistically significant, and suppressed mesenchymal markers expression of xenograft. The mesenchymal markers expression of N-cadherin, Vimentin, α-SMA, SMAD3, p-SMAD3 and endothelial markers E-cadherin, VE-cadherinin in xenografts were determined.

## Discussion

Through EMT (epithelial-mesenchymal transition), a mesenchymal phenotype in epithelial cells can be acquire, wherein the marker expression and acquired functions of mesenchymal cell, in addition to lost the epithelial functions and markers expression of epithelia. Besides, migration and delamination of epithelial cells can be caused by EMT, then mesenchymal cells derived from epithelial cell can migrate into the underlying tissue [36]. The ligands of TGF-β (transforming growth factor β) family often initiate the cellular process, EMT [37]. Many investigations aim at the markers of EMT from thirty-two cancer types [38, 39] and the roles of TGF-β pathway [38, 40]. A number of elements are important factors in regulation of gene expression and EMT, while referred to development of TNBC [41-44].

Moreover, the transcriptional regulation mechanism of EMT and how miRNA modulated EMT or development of TNBC was unclear yet. Recent research revealed that miR-34 might serve as inhibitor in Nasopharyngeal carcinoma (NPC) by targeting TNBC-derived growth factor (HDGF). It indicated that miR-34 acted as a tumor suppressor in NPC, and may be a novel therapeutic target for the treatments of patients with NPC [45]. Moreover, over-expression of miR-425 is induced by IL-1β, which aim at 3’UTR of miR-425, could regulate homolog expression of tensin and phosphatase. The activation of NF-κB induced by IL-1β up-regulates miR-425 expression, while enhances transcription of miR-425 gene. Therefore, the proliferation of gastric cancer cell is promoted by miR-425 through suppressing tensin homolog and phosphatase. In summary, it confirm that over-expression of miR-425 dependent on NF-κB plays the critical role for the above process [46]. EMT could induce extracellular cytokine TNF-α, TGF-β, PDGF, HGF or bFGF to bond with its receptor in BC, and then activate intracellular effectors, such as Ras, NF-κB, Smads, etc. Therefore, in this study, miRNAs were ascertained and searched after its effects on EMT and development process of TNBC.

At first, the differential miRNAs expression profiles between tumor and para-tumor tissues were assessed with miRNA-sequencing experiments. Then, we found that, compared with para-tumor tissues, 9 miRNAs were upregulated and 3 miRNAs (included miR-425) were down-regulated in tumor tissues of patients with TNBC. Furthermore, the relative expression level of miR-425 in non-tumor, para-tumor, and tumor tissues were determined using qRT-PCR method. This work identified relative levels of miR-425 expression in tumor tissues were obviously less than that in para-tumor and non-tumor tissues. However, there was no obvious difference between para-tumor and non-tumor tissues. Consequently, the relative miR-425 expression levels were investigated in two cell lines of human TNBC. The results confirmed that miR-425 expression level was significantly decreased in the basal epithelial phenotype cell lines Ts 578T and mesenchymal phenotypic cell line MDA-MB-231, which indicatd that miR-425 down-expression was related to phenotype of EMT. Therefore, the targets of miR-425 were foretasted with Targetscan, an online software, and TGF-β1 was found that it is the possibly target of 3’ UTRs of TGF-β1 mRNA. Then, further experiments were performed and identified that miR-425 could straight bind with 3’ UTRs of TGF-β1 mRNA. In a word, miR-425 directly targeted to TGF-β1 mRNAs. Hence, miR-425 could straight control mRNA and the protein expression level of TGF-β1, and furtherly affect expression levels of E-cadherin, VE-cadherin, N-cadherin, Vimentin, α-SMA, and SMAD3 protein in TNBC cell lines. Taken together, our study maybe point out a novel aim at treatment for TNBC associated with EMT.

Besides, to assess the biofunctions of miR-425, several molecular functional tests were performed in TNBC cell lines transfected with mimics or inhibitors of miR-425, respectively. It identified that miR-425 suppressed proliferation process and the migration or invasion ability of TNBC cell lines, and significantly promoted the expressions of epithelial markers E-cadherin and VE-cadherin, and suppressed mesenchymal markers expression of N-cadherin, Vimentin, α-SMA, total and phosphorylated SMAD3 (p-SMAD 3) in TNBC cell lines. Our results demonstrated that miR-425 targets to TGF-β 1, and was a crucial mediator of EMT and development of TNBC through regulating TGF-β1/SMAD3 signaling pathway. Moreover, we did not found that it is involved in well-known molecules such as mTOR, Ki-67, etc.

Furthermore, the effects on Human TNBC xenografts and its mechanism were investigated using miR-425 agomir. Our results conirmed that the xenografts of control groups grew rapidly. But in the experiment group, which treated with agomir of miR-425, the xenografts grew slowly *in vivo*. Moreover, the size and weight of tumors in experiment group (treated with agomir of miR-425) was significantly less than that of control group. At the time of end, the average tumor volume of control group was much larger than that of experiment group, and the difference between the two groups had statistical significance. Therefore, the agomir of miR-425 could obviously inhibit tumor proliferation and its development process. Moreover, treatment with agomir of miR-425 could down-regulate the expression of TGF-β1 and SMAD3 with a statistically significant, while suppressed the expression of mesenchymal cell markers and promoted epithelial markers E-cadherin and VE-cadherin expression in xenografts.

We thought that our manuscript was obviously composed only one part of the whole. The assays *in vivo* and *in vitro* were all separately used to identify our conclusions. In summary, we illustrated that miR-425 targets to TGF-β1. Moreover, miR-425 was a crucial suppressor of EMT and the development of TNBC through inhibiting TGF-β1/SMAD3 signaling pathway. It suggested that aim at TGF-β1/SMAD3 signaling pathway through enhancing miR-425 expression, was a feasible therapy strategy for TNBC.

## Acknowledgments

Not applicable.

## Availability of data and materials

The datasets used and/or analyzed during the current study are available from the corresponding author on reasonable request.

## Authors' contributions

LYP and ZY conducted the cell and animal experiments and collected the data and wrote the manuscript. LYP, YF, and CJL analyzed and interpreted the data from the experimental results. CJL and ZJQ were involved in the conception and design of the study, and revised the manuscript. All authors have read and approved the final version of the manuscript.

## Ethics approval and consent to participate

All experiments related to animals were performed based on Laboratory Animal Center, Capital Medical University, Beijing, China, Ethical Committee Acts. The experimental mice were dealt with according to the standards supported by the Animal Protection Committee of Capital Medical University. Moreover, the ethical approval was obtained for the animal experiments conducted in your study from Animal Protection Committee of Capital Medical University.

